# Population bottlenecks shape laboratory evolution of piperacillin-tazobactam resistance in *Klebsiella grimontii* and reveal a shared within-patient evolutionary trajectory

**DOI:** 10.64898/2026.06.15.732307

**Authors:** Ellie Allman, Aakash Khanijau, Rachel Mcgalliard, Richard N. Goodman, Christopher M. Parry, Enitan D. Carrol, Nicholas A. Feasey, Fabrice E. Graf, Adam P. Roberts

## Abstract

Laboratory-based experimental evolution is widely used to investigate how antimicrobial resistance (AMR) emerges and to identify resistance-associated trade-offs that could inform treatment strategies. However, there is limited understanding of how *in vitro* AMR evolution reflects the complexity of resistance evolution within the human host, where selective pressures, and therefore evolutionary pathways, are more variable. Here, we investigated the effect of population bottleneck size and growth environment on the evolution of piperacillin-tazobactam (TZP) resistance in *Klebsiella grimontii* and compared this to resistance evolution observed during a recurrent bloodstream infection. Three clonal *K. grimontii* isolates cultured from one patient over four months included a TZP-susceptible ancestor and a within-patient evolved TZP-resistant isolate. The susceptible ancestor was evolved under TZP selection using either a small 0.1% bottleneck or a larger 5% bottleneck, and under a second environment, LB supplemented with 5% sheep blood, using a 0.1% bottleneck. Evolved isolates were assessed for TZP susceptibility, β-lactamase activity, fitness, and genomic changes. A single nucleotide polymorphism (SNP) in the promoter region of the chromosomally located β-lactamase gene *bla*_OXY-6-4_ was identified in the within-patient evolved isolate and was replicated in all 0.1% bottleneck lineages across both environments. In contrast, the larger 5% bottleneck lineages exhibited greater phenotypic variation and genetic diversity, including multiple *bla*_OXY-6-4_ promoter variants and variable TZP MICs. These findings show that laboratory evolution can reproduce key within-patient resistance mechanisms, but that bottleneck size strongly shapes the resistance phenotypes and mutational landscapes observed *in vitro*.

**Importance:** Adaptive laboratory evolution is increasingly used to predict how antimicrobial resistance emerges and to identify trade-offs associated with resistance acquisition that could inform future treatment strategies. Here, we directly compared piperacillin-tazobactam resistance evolution in the laboratory with resistance that emerged within a patient during a recurrent bloodstream infection. We show that a small population bottleneck reproducibly selected the same *bla*_OXY-6-4_ promoter mutation observed in the patient, whereas a larger bottleneck produced more diverse evolutionary outcomes. These findings build on previous work showing that experimental conditions shape laboratory evolution outcomes and highlight population bottleneck size as an important experimental parameter when designing laboratory evolution studies that intend to model clinically relevant resistance evolution.

## Introduction

Antimicrobial resistance (AMR) remains a major global health concern, and improved understanding of resistance emergence and persistence is essential for developing strategies to combat it. While many studies have explored the molecular mechanisms underpinning resistance, our ability to predict how bacteria evolve under different selective pressures and growth environments remains limited. Insights into the evolutionary pathways favoured following antibiotic exposure could inform the development of more effective, evolution-informed antimicrobial treatment strategies(1). Experimental evolution therefore offers a potentially powerful tool to study the dynamics of resistance development under controlled conditions(2).

Resistance acquisition is often associated with evolutionary trade-offs, such as reduced fitness in the absence of antibiotics, or increased susceptibility to alternative drugs, known as collateral sensitivity(3–5). These trade-offs have generated interest within therapeutic strategies, including antibiotic cycling or sequential therapy, to slow resistance evolution(6). However, the reproducibility and clinical relevance of experimental evolution approaches in understanding real-world AMR dynamics are uncertain, particularly as most experimental evolution studies are performed under single defined laboratory conditions(3, 7–11).

A key question is whether *in vitro* experimentation accurately reflects the selective pressures and evolutionary trajectories that occur within human hosts. Experimental conditions, including pH, temperature, and nutrient sources, can influence resistance evolution and associated trade-offs(6, 12, 13). Host-like conditions and population bottlenecks have also been highlighted as important factors shaping evolutionary trajectories, but in the real-world, these are incredibly diverse and difficult to represent fully in model systems(14). Few studies have examined how *Klebsiella* spp. evolve under host-relevant conditions, or whether selective dynamics seen *in vitro* reflect those occurring during clinical infection, despite *Klebsiella* spp. being a World Health Organisation AMR priority pathogen.

*Klebsiella grimontii* are Gram-negative bacteria belonging to the order Enterobacterales. It is an emerging pathogen associated with wound infections, bacteraemia, and antibiotic-associated haemorrhagic colitis. *K. grimontii* is commonly misidentified as *Klebsiella oxytoca*, but can be distinguished by *gyrA*, *rpoB*, and the chromosomally encoded class A β-lactamase, *bla*_OXY-6_(15). While *bla*_OXY-6_ typically confers low-level β-lactamase activity, promoter mutations can increase expression leading to resistance to penicillins and other β-lactams, including narrow-spectrum cephalosporins and aztreonam(16–18). Due to frequent misidentification of *K. grimontii*, limited data exist for the antimicrobial treatment and resistance patterns of this species(15). Typically, treatment for *Klebsiella* spp. infections is the same as antibiotic therapy for other Enterobacterales species(16). However, increasing prevalence of extended-spectrum β-lactamase (ESBL)-producing Gram-negative bacteria has contributed to increased use of β-lactam/β-lactamase inhibitor combinations, such as piperacillin-tazobactam (TZP), in England(19, 20).

In this study, we describe TZP resistance evolution in *K. grimontii* from a patient with a recurrent bloodstream infection which showed increased resistance over four months. Our aim was to assess how environmental context and population bottleneck size influence evolutionary trajectories of TZP resistance, and to evaluate the extent to which *in vitro* evolution mirrors within-host resistance evolution.

## Results

### Within-patient TZP resistance evolution of linked clonal *Klebsiella grimontii* isolates

VS1520 was isolated at the onset of the initial bloodstream infection episode, and VS1543 isolated four days later. Both were identified as *K. grimontii* and were TZP-susceptible with identical resistance patterns revealed by routine susceptibility testing. The patient was initially prescribed TZP for nineteen days to treat the infection. One hundred and thirty days after the initial infection episode, MB2616 was isolated from an additional infection episode and identified as TZP-resistant, indicating that TZP resistance had likely evolved within the patient. Between the first and third isolate being cultured, the patient was empirically treated with several different antimicrobials, including two further rounds of TZP treatment as seen in Fig. 1. Clonality of these isolates was supported by comparing SNPs and indels between isolates. We identified one insertion between VS1520 and VS1543, and 13 SNPs and one insertion between VS1520 and MB2616 using Breseq. Additionally, VS1543 and MB2616 both showed missing coverage across unresolved regions of a Col(pHAD28) plasmid. Average nucleotide identity (ANI) analysis of hybrid assembled genomes using pyani (ANIm) showed ANI values of > 99.99% between all three isolates. Species identification and multi-locus sequence type of all three clinical isolates was predicted as *K. grimontii* ST362 using Pathogenwatch. Together, these findings indicate that the three clinical isolates are derived from a common ancestor.

**Figure 1:**
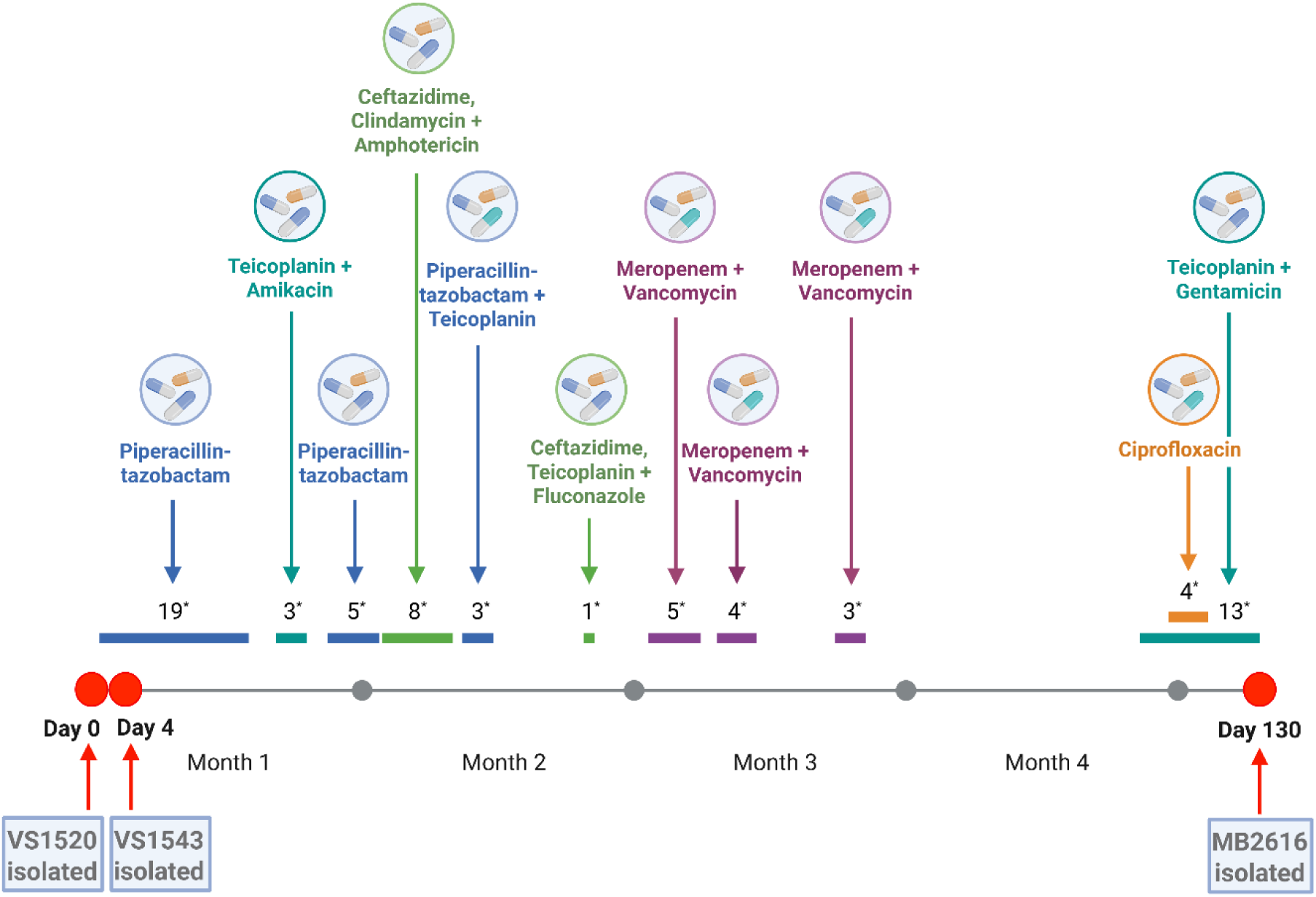
Patient antimicrobial treatment timeline. Timeline of antimicrobial therapy from day 0 to day 130 of hospitalisation. Red markers indicate timepoints at which *K. grimontii* isolates were cultured from blood samples. Asterisks denote number of days each treatment was given. Image created using BioRender.

#### *bla*_OXY-6-4_ promoter mutation leads to **β-**lactamase hyperproduction and TZP resistance phenotype

We tested the minimum inhibitory concentrations (MIC) of the three of clinical isolates and confirmed susceptibility to TZP (1/4 µg/ml) for VS1520 and VS1543, and high-level TZP resistance (128/4 µg/ml) for MB2616. Additionally, MB2616 showed resistance to other β-lactam antibiotics, including amoxicillin/clavulanic acid, 2^nd^ generation cephalosporins, cefuroxime and cefoxitin, and 3^rd^ generation cephalosporin, cefpodoxime. Intermediate resistance to aztreonam was observed and MB2616 was also resistant to ciprofloxacin and colistin but susceptible to meropenem (Table 1). To investigate the molecular basis of β-lactam resistance, we analysed the genomes of the *K. grimontii* clinical isolates using ResFinder and identified three genes inferred in resistance located on the chromosome in all three isolates; *bla*_OXY-6-4_ β-lactamase gene, and *oqxA*- *oqxB* efflux pump encoding genes. There was no difference in the coding sequences of *bla*_OXY-6-4_ between the isolates. However, the promoter region contained a single nucleotide polymorphism (SNP) in the −10 promoter element of MB2616 (Supplementary Fig. 1). This G→A transition, located 33 bp upstream of the start codon and corresponding to the fifth base of the −10 promoter element, may increase *bla*_OXY-6-4_ expression and be responsible for the observed resistance phenotype. This SNP has previously been reported in *bla*_OXY_ promoters of the *K. oxytoca* complex(16, 18). To test this, we quantified the β-lactamase activity using a nitrocefin assay and confirmed that the TZP-resistant isolate hyperproduced a β-lactamase compared to TZP-susceptible VS1520 and VS1543 (Supplementary Fig. 2). Nitrocefin hydrolysis was significantly higher in MB2616 (0.93 ± 0.016) compared to VS1520 (0.51 ± 0.035, *p* < 0.0001), and VS1543 (0.47 ± 0.017, *p* < 0.0001). No significant difference was observed between VS1520 and VS1543 (*p* = 0.4666).

**Table 1:**
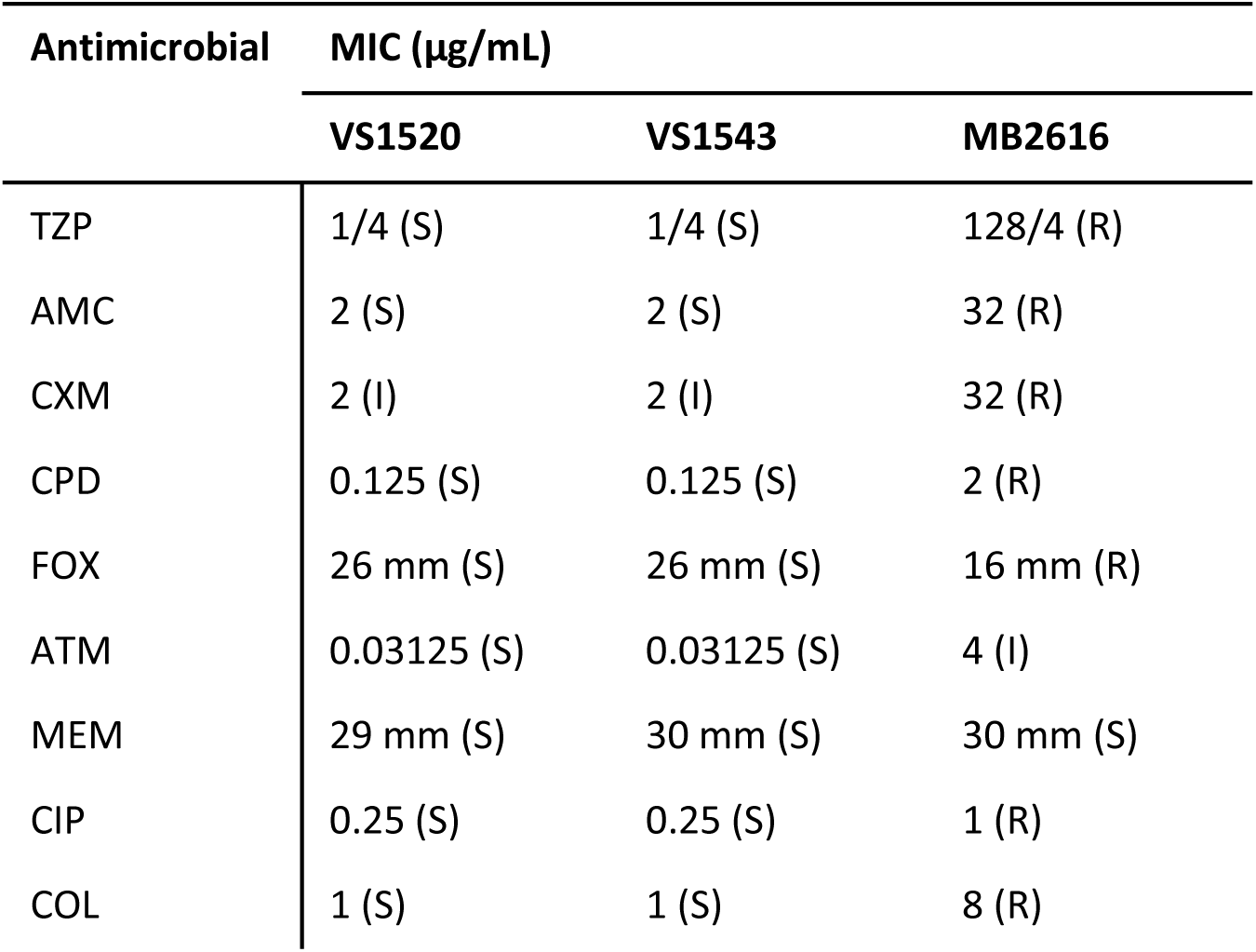
Antimicrobial susceptibility profiles of *K. grimontii* isolates VS1520, VS1543, and MB2616. Minimum inhibitory concentrations (MICs; µg/mL) or disk diffusion zone diameters (mm) are shown for piperacillin/tazobactam (TZP), amoxicillin/clavulanic acid (AMC), cefuroxime (CXM), cefpodoxime (CPD), cefoxitin (FOX), aztreonam (ATM), meropenem (MEM), ciprofloxacin (CIP), and colistin (COL). Interpretation of susceptibility is based on EUCAST guidelines (S = susceptible, I = susceptible with increased exposure, R = resistant). Antimicrobial susceptibility testing was performed in triplicate (*n* = 3).

#### Experimental conditions change the outcome of *in vitro* evolution to TZP resistance

We evolved the TZP susceptible ancestor (VS1520) in sub-inhibitory concentrations of TZP in LB broth, followed by selection on TZP-supplemented LB agar. We compared two bottleneck sizes; 0.1% and 5% passage volumes, which allowed for a direct comparison of how bottleneck size influences evolutionary outcomes in a standard LB environment. An independent evolution experiment was performed in LB supplemented with 5% sheep blood using a 0.1% bottleneck.

Evolution of TZP resistance was achieved in all 0.1% bottleneck lineages, regardless of growth environment (Fig. 2). The MIC for TZP varied across biological replicates in most of the *in vitro* evolved isolates, reflecting phenotypic heterogeneity within the evolved populations. In contrast, the within-patient evolved MB2616 showed a consistent TZP MIC of 128/4 µg/mL. Greater phenotypic variation was observed in the 5% bottleneck lineages, with some evolved lineages (*n* = 6) exhibiting TZP susceptibility, despite initially growing on TZP-supplemented agar. All antibiotic-free control isolates (*n* = 9) across all three conditions remained TZP-susceptible (MIC = 1/4 µg/mL) (Supplementary Table 2).

**Figure 2:**
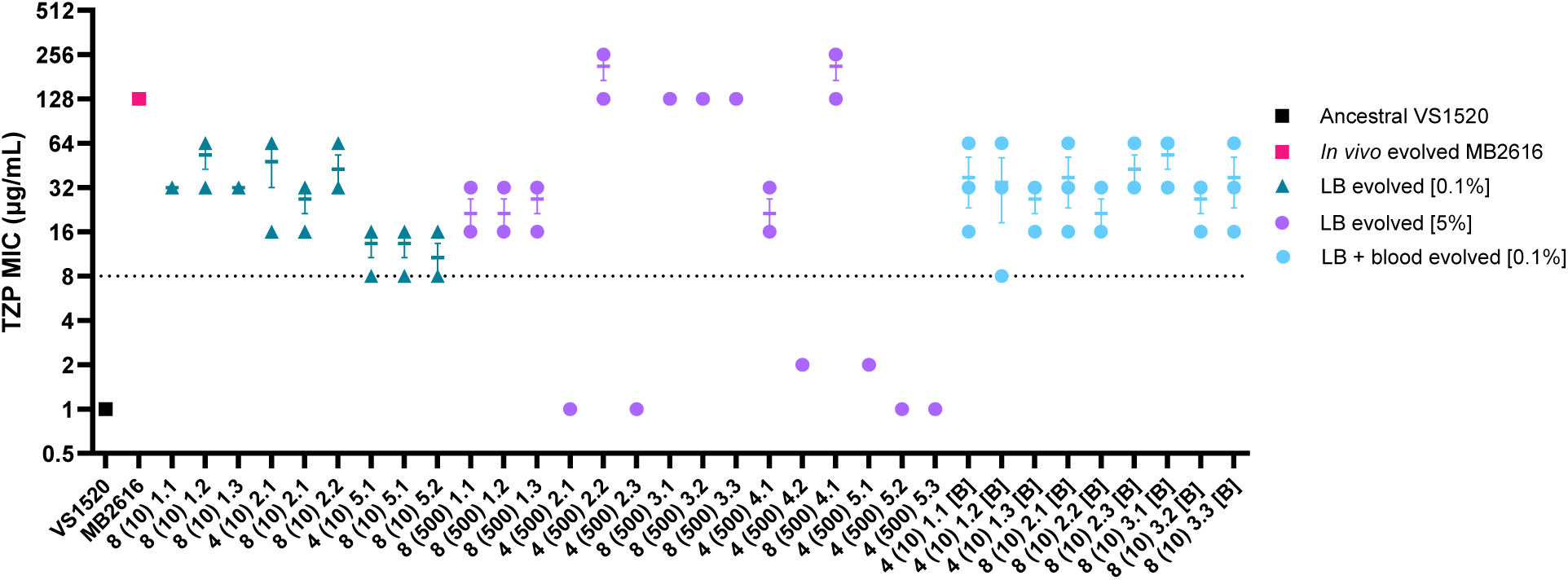
Minimum inhibitory concentration (MIC) of clinical isolates and *in vitro* evolved isolates against piperacillin-tazobactam (TZP). Tazobactam concentration was fixed at 4 µg/mL. Antimicrobial susceptibility testing was performed in triplicate (n = 3). Error bars represent mean ± SEM.

#### Increased β-lactamase activity in TZP evolved isolates

In the LB (0.1%) condition (Fig. 3A), all TZP-evolved isolates exhibited significantly increased β-lactamase activity compared to the ancestor (*p* < 0.0001) and all antibiotic-free controls showed activity comparable to the ancestor (*p* ≥ 0.8895), confirming that hyperproduction was selected for by TZP exposure.

**Figure 3:**
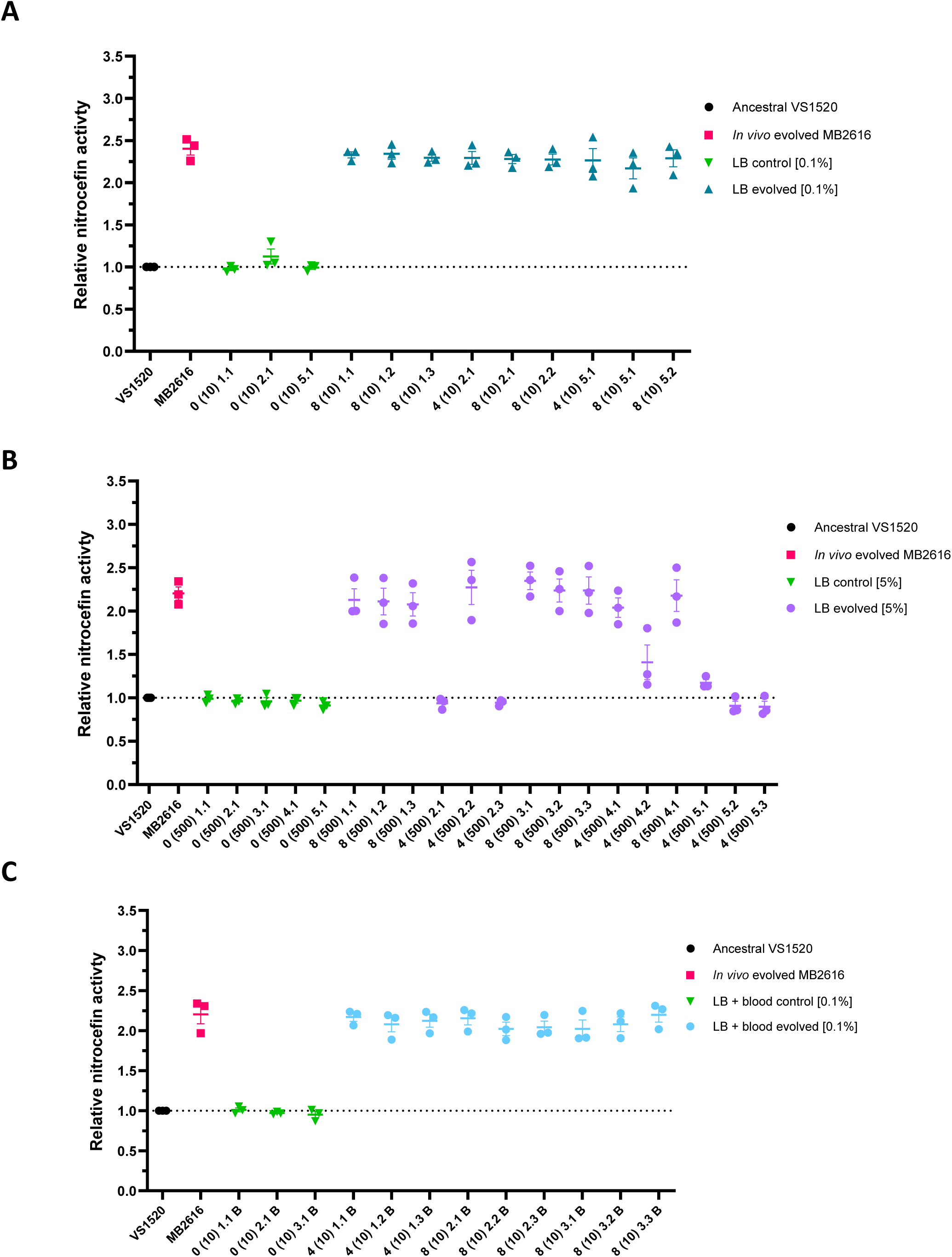
Relative nitrocefin activity of patient evolved MB2616 and TZP-evolved isolates compared to ancestral VS1520. Hydrolysis of the chromogenic β-lactam substrate, nitrocefin (50 µM), was quantified over 30 min by measuring absorbance at 486 nm every 90 s and measuring the area under the curve (AUC) relative to the susceptible ancestor. Error bars represent standard error of the mean (*n* = 3). **A,** LB + TZP (0.1%) evolved lineages. **B,** LB + TZP (5%) evolved lineages. **C,** LB + blood + TZP (0.1%) evolved lineages.

The LB (5%) condition (Fig. 3B) showed substantial variability of nitrocefin hydrolysis, consistent with variability in susceptibility profiles (Fig 2). Several isolates showed increased hydrolysis (*p* < 0.0001), correlating with TZP MICs above 16/4 µg/mL. Other isolates, particularly those with MICs ≤ 2/4 µg/mL, did not differ significantly from the ancestor (*p* ≥ 0.1172). Antibiotic-free controls exhibited no significant change in activity (*p* > 0.9999).

In the LB + blood (0.1%) condition (Fig. 3C), all TZP-evolved isolates consistently hyperproduced β-lactamase (*p* < 0.0001), mirroring the LB (0.1%) pattern. Control isolates again showed no significant difference from the ancestor (*p* > 0.9999). All *p*-values and mean differences are reported in Supplementary Table 3.

### Promoter SNP in *bla*_OXY-6-4_ reproduced in *in vitro* experimental evolution with small bottleneck

We compared the *K. grimontii bla*_OXY-6-4_ β-lactamase gene promoter sequences of within-patient evolved and the laboratory-evolved strains (Fig. 4) and showed the *bla*_OXY-6-4_ promoter SNP (G→A transition in the fifth base of the −10 sequence), observed in the within-patient evolved isolate MB2616, was replicated in all 0.1% bottleneck mutants across both environments (LB and LB + 5% sheep blood).

**Figure 4:**
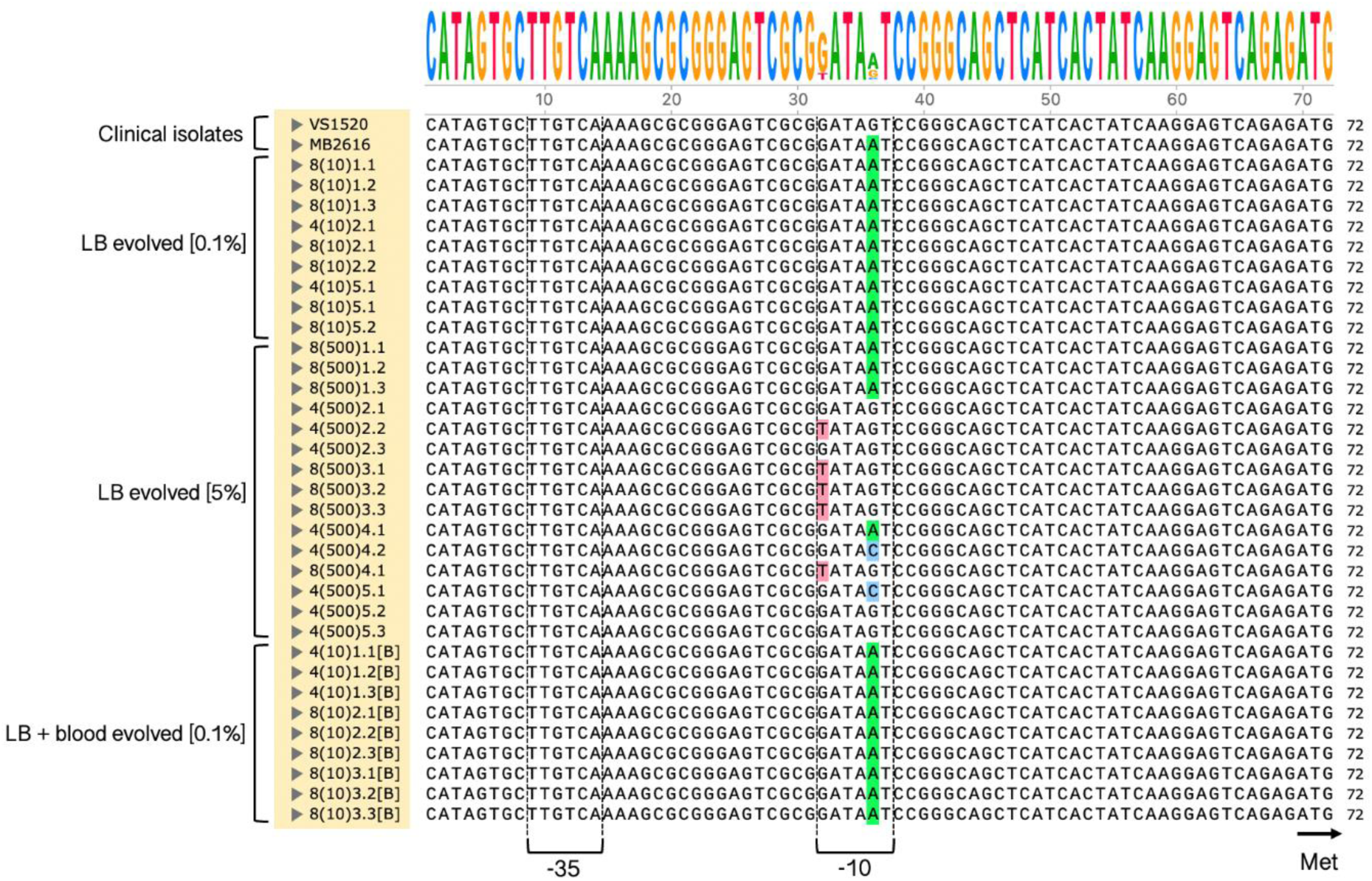
Multiple sequence alignment of *K. grimontii bla*_OXY-6-4_ β-lactamase gene promoter sequences in VS1520, MB2616, and laboratory-evolved lineages across all conditions. The −10 and −35 promoter elements are labelled, and the start codon is marked Met. Highlighted bases are colour coded and represent SNPs relative to the ancestral isolate.

In contrast, the larger 5% bottleneck lineages exhibited greater genetic diversity in *bla*_OXY-6-4_ promoter sequences, with distinct mutations correlating with a range of phenotypic resistance levels. Of the 15 isolates in this group, four retained the wild-type (WT) promoter sequence and displayed TZP MICs of 1/4 µg/mL, indicating that resistance did not evolve in these isolates. In the remaining isolates, four had the G→A transition in the fifth base of the −10 sequence as seen in MB2616, producing an MIC against TZP between 16/4 – 32/4 µg/mL. Two isolates exhibited a G→C transversion in the fifth base of the −10 sequence, which slightly increased the MIC to 2/4 µg/mL but did not confer a TZP-resistant phenotype. The remaining five isolates from the LB (5%) group displayed a G→T transversion in the first base of the −10 region. These isolates demonstrated the highest TZP MIC, ranging between 128/4 and 256/4 µg/mL, highlighting the significant impact of this mutation on β-lactamase expression.

#### Genomic comparison of the susceptible ancestor to within-patient and *in vitro*-evolved isolates

We compared the hybrid assembled genome of the ancestral isolate, VS1520, to short-read genomes of the two within-patient evolved isolates to identify mutations that accumulated over the course of infection (Fig. 5). Trimmed reads were aligned to the VS1520 genome using Breseq to identify SNPs and indels. All three isolates harboured an IncFIB(K) plasmid with 100% identity, and a Col(pHAD28) plasmid which was not identical between isolates. No acquired AMR genes were located on either plasmid. Comparison of the two TZP-susceptible isolates, VS1520 and VS1543, revealed only a single mutation: a 1 bp insertion in the xanthine-guanine phosphoribosyltransferase gene (*gpt*), which is involved in the purine salvage pathway(21). VS1543 also showed missing read coverage across regions corresponding to the Col(pHAD28) plasmid, likely due to assembly artefacts in repetitive plasmid regions.

**Figure 5:**
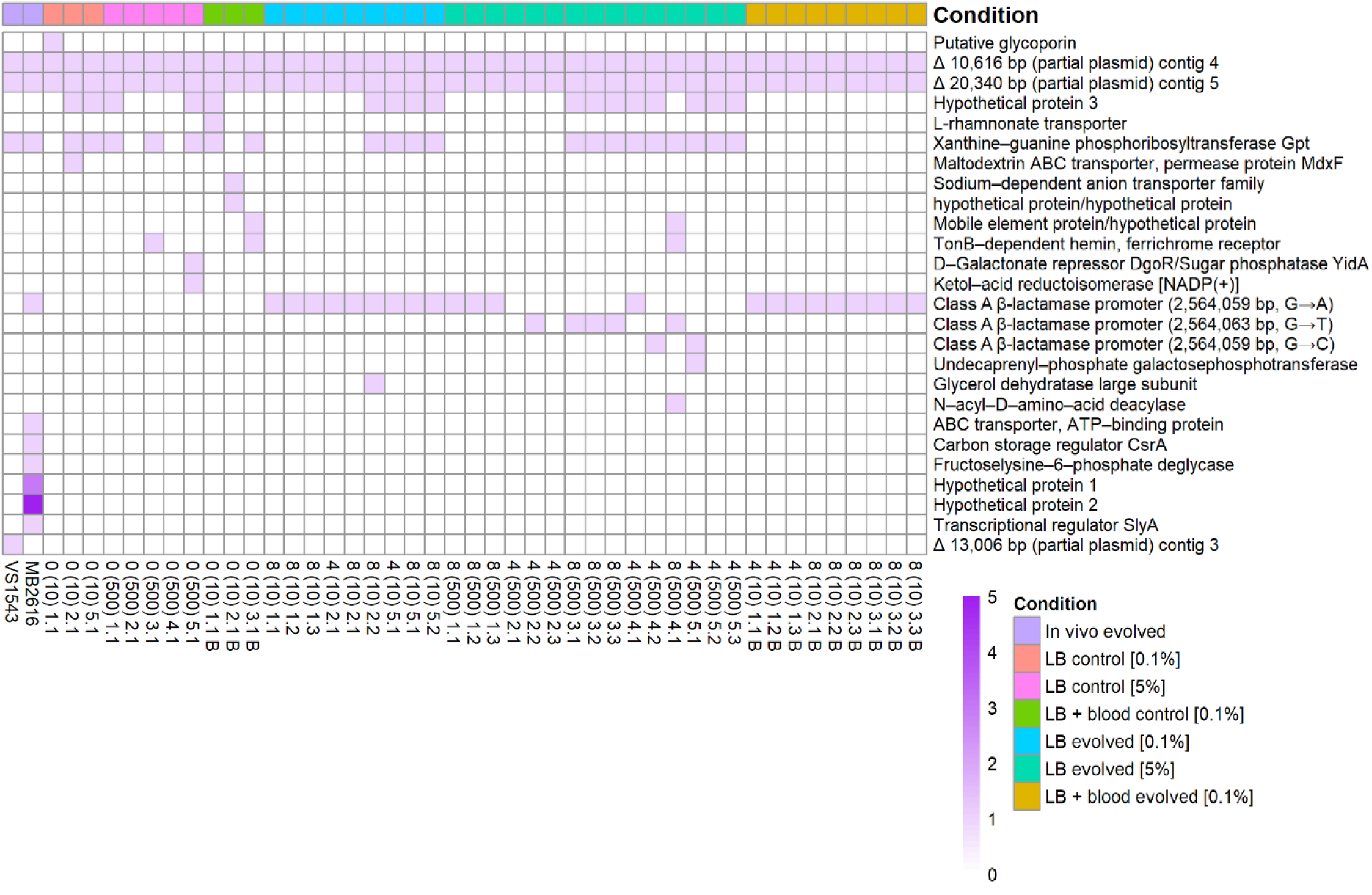
Heatmap showing SNPs and indels in evolved lineages relative to the ancestral strain, VS1520, under different experimental conditions. Using Breseq, the hybrid assembled genome of ancestral VS1520 was compared to paired-end short reads of within-patient evolved and laboratory-evolved isolates to identify SNPs and indels accumulated. Regions marked with ‘Δ’ denote deletions, and colour intensity corresponds to the number of mutations detected in each gene.

When comparing VS1520 to resistant MB2616, we identified 13 SNPs and one insertion across eight different genes, along with missing read coverage across two regions of the Col(pHAD28) plasmid, consistent with the patterns described above. This included a promoter SNP in *bla*_OXY-6-4_ (G→A in the - 10 element), as shown above (Fig. 4). Additional mutations included a 1 bp insertion in the *gpt* gene, as also seen in VS1543. A missense substitution was identified in a gene encoding the ATP-binding subunit of an ATP-binding cassette transporter, which hydrolyses ATP and provides energy for substrate transport across the membrane in processes such as nutrient uptake, toxin secretion, and antibiotic efflux(22, 23). Another missense substitution was observed in the carbon storage regulator *csrA*, which has a role in the regulation of multiple metabolic pathways including motility, glycogen synthesis, adherence, and biofilm formation(24). Additionally, a nonsense substitution was observed in *slyA*, a transcriptional regulator required for capsule formation and regulation of virulence-associated genes(25, 26). Furthermore, a nonsynonymous substitution occurred in the gene encoding fructoselysine-6-phosphate deglycase. This enzyme is involved in catalysing the conversion of fructoselysine 6-phosphate into glucose 6-phosphate and lysine(27). Together, these mutations suggest that MB2616 accumulated multiple genetic changes during *in vivo* adaptation.

We then investigated mutations acquired in laboratory-evolved mutants relative to the ancestor VS1520 (Fig. 5). All isolates retained both plasmids, and consistent regions of missing read coverage were observed within the Col(pHAD28) plasmid across all isolates (denoted as ‘Δ 10,616 bp (partial plasmid) contig 4’ and ‘Δ 20,340 bp (partial plasmid) contig 5’), in line with the assembly artefacts described above.

All LB (0.1%) mutants carried the *bla*_OXY-6-4_ promoter SNP (G→A), as shown in Fig. 4. Four isolates also shared a 1 bp insertion (+ T) in *gpt* and a 1 bp insertion (+ G) in hypothetical protein 3. One isolate had a 1 bp deletion in the glycerol dehydratase large subunit gene. The antibiotic-free evolved controls (*n* = 3) exhibited a range of SNPs/indels despite undergoing no antibiotic selection, including the same insertions in hypothetical protein 3 and *gpt*.

In the larger bottleneck condition (LB (5%)), greater genetic diversity was seen. Multiple *bla*_OXY-6-4_ promoter variants were present (Fig. 4), and four isolates lacked promoter SNPs and remained susceptible to the level of the ancestor (Fig 2 and 3B). Nine mutants exhibited an insertion (+ T) in *gpt*, of which eight also carried a 1 bp insertion (+ G) in hypothetical protein 3, as seen across the 0.1% bottleneck condition. One isolate (8 (500) 4.1) showed multiple unique mutations: in addition to its *gpt* insertion, missense substitutions were also identified in genes encoding N-acyl-D-amino-acid deacylase and TonB-dependent hemin ferrichrome receptor. A 4 bp deletion was found in an intergenic region (mobile element protein/hypothetical protein) on the IncFIB(K) plasmid. Another isolate (4 (500) 5.1) had a 1 bp deletion in undecaprenyl–phosphate galactose phosphotransferase. Antibiotic-free controls (*n* = 5) also acquired diverse SNPs and indels, including *gpt* and hypothetical protein 3 insertions.

In the LB + Blood (0.1% bottleneck) condition, all TZP-evolved mutants carried the *bla*_OXY-6-4_ promoter SNP (G→A), as shown in Fig. 4. No additional background mutations were observed, aside from the plasmid-associated coverage differences noted across all lineages. In contrast, antibiotic-free controls acquired varied mutations, including insertions in *gpt* and hypothetical protein 3, a deletion in the L-rhamnonate transporter, and SNPs in a sodium-dependent anion transporter and a TonB-dependent hemin/ferrichrome receptor (Fig. 5).

These findings support clonal relatedness between the clinical isolates and show that the *bla*_OXY-6-4_ promoter mutation (G→A) was reproducibly selected under TZP pressure in all small bottleneck (0.1%) conditions, regardless of growth environment. Although some shared variants were observed between within-patient and *in vitro*-evolved lineages, including in *gpt*, the recurrent insertions in *gpt* and hypothetical protein 3 were also detected in antibiotic-free controls. Additionally, the *gpt* insertion was present in the ancestral long-read only assembly but absent from the short-read and hybrid assemblies. MB2616 accumulated additional mutations linked to clinical adaptation that were absent from *in vitro*-evolved lineages. Moreover, the diversity observed in antibiotic-free controls highlights background adaptation unrelated to antibiotic selection and helps distinguish background changes from mutations likely associated with resistance evolution.

#### Laboratory evolution of TZP resistance does not incur consistent fitness costs or collateral trade-offs

Relative fitness was measured by growth compared to the ancestor’s growth in liquid media in the same environment in which each group evolved: LB broth for isolates from the LB (0.1%) and LB (5%) evolved lineages, and LB supplemented with 5% sheep blood for isolates from the LB + blood (0.1%) condition. No significant differences in relative fitness were observed between the ancestor and lab-evolved isolates from the LB (0.1%) and LB (5%) conditions (*p* > 0.05 for all comparisons)., whereas the within-patient evolved isolate MB2616 exhibited a measurable fitness cost (*p* ≤ 0.0021) (Supplementary Fig. 3A and Fig. 3B). In the LB + blood (0.1%) condition, MB2616 again showed reduced fitness (*p* = 0.0002), and some TZP-evolved isolates showed reduced fitness compared to VS1520 (*p* ≤ 0.0079), although high variation was observed likely caused by haemolysis of the blood (Supplementary Fig. 3C).

To investigate potential collateral effects, representative TZP-evolved isolates were selected based on their mutational profiles to capture unique genomic backgrounds (*n* = 8). Susceptibility testing revealed no consistent collateral sensitivity patterns across the clinically relevant antibiotics tested (colistin, gentamicin, ciprofloxacin, meropenem, tigecycline, and trimethoprim-sulfamethoxazole). A slight, non-significant two-fold reduction in colistin MIC was observed across the tested TZP-evolved isolates relative to the ancestor, but this trend was not consistent across all replicates (Supplementary Fig. 4). While recent work suggests that β-lactamase expression can lead to collateral sensitivity to colistin in Enterobacterales, this effect was not robustly observed here(28). Together, these findings indicate that TZP resistance evolution *in vitro* did not incur any clinically useful evolutionary trade-offs.

## Discussion

In this study we investigated the effect of bottleneck size and growth environment on the evolution of TZP resistance in *K. grimontii* and compared this to a within-patient evolved isolate that developed TZP resistance during infection.

Two distinct pathways to TZP resistance were identified *in vitro*: a G→T transversion in the first base, and a G→A transition in the fifth base of the −10 promoter sequence of *bla*_OXY-6-4_. The G→A SNP present in the within-patient evolved isolate was reproducibly selected in all *in vitro* lineages under the smaller 0.1% bottleneck, regardless of growth condition (LB or LB + 5% sheep blood). This SNP, located in a critical regulatory region of the promoter, was associated with β-lactamase hyperproduction and elevated TZP MICs, ranging between 8/4 – 64/4 µg/mL. Both promoter mutations have been previously described in *bla*_OXY_ and are associated with differing increases in promoter strength, with the G→T transversion in the first base of the −10 element conferring a stronger effect than the G→A transition in the fifth base(29, 30). Given that within-host bottlenecks can be extreme and sometimes even single-cell(31, 32), selection of this SNP under a stringent population bottleneck suggests that it is a highly adaptive mutation that emerges under strong selective pressure. Its reproducible emergence both *in vitro* and within-patient highlights convergence between experimental and within-patient resistance pathways, an important consideration if laboratory evolution is to be used to inform treatment strategies. However, *in vitro*-evolved isolates carrying the same G→A promoter mutation as MB2616 showed a range of TZP MICs, suggesting that additional within-patient mutations, together with differences in fitness and genetic background, may influence the final resistance phenotype.

The larger 5% bottleneck lineages demonstrated greater genetic heterogeneity, both in *bla*_OXY-6-4_ promoter mutations and across the wider genome. This illustrates the interplay between population bottleneck size, selection, and mutation fixation. Under the smaller bottleneck, the G→A SNP was consistently selected, whereas the larger bottleneck allowed multiple promoter variants to emerge, with varying effects on β-lactamase activity (Fig. 3) and TZP resistance (Fig. 2), including a G→C substitution that only slightly increased TZP MIC. The presence of isolates that remained phenotypically susceptible despite growth on TZP-supplemented agar suggests increased phenotypic heterogeneity within the larger bottleneck populations. This may be due to mechanisms such as cross-protection from neighbouring resistant cells, or transient antibiotic tolerance. This is consistent with the larger effective population sizes in the 5% bottleneck condition, where less severe bottlenecks allow multiple variants to be retained across passages. In contrast, the reduced diversity observed under the 0.1% bottleneck reflects a smaller effective population size, increasing the impact of genetic drift and limiting the retention of genetic variation.

Despite the reproducibility of the *bla*_OXY-6-4_ promoter mutation under the small bottleneck, most additional mutations identified in the within-patient evolved isolate were not observed in laboratory-evolved strains. Although *gpt* variation was seen in both within-patient and *in vitro*-evolved isolates, its significance remains unclear because the insertion was also detected in antibiotic-free controls. A similar pattern was seen for the insertion in hypothetical protein 3, which frequently co-occurred with the *gpt* insertion in laboratory-evolved isolates but was absent from the within-patient evolved isolates. Together, this suggests that these insertions may reflect variation in the founding ancestral population and/or background changes arising during growth in laboratory media, rather than mutations specifically associated with TZP exposure. This is further supported by comparison of the ancestral *gpt* sequence across the short-read, hybrid, and long-read only assemblies, where the insertion was present only in the long-read assembly. As these datasets were generated from different ancestral colonies, this suggests that the founding ancestral population may not have been homogeneous.

The remaining mutations in MB2616 included missense and nonsense substitutions in genes involved in metabolism, nutrient acquisition, virulence regulation, and membrane transport. These mutations may have facilitated adaptation and persistence within the host, suggesting that the broader mutational landscape is shaped by additional host-specific pressures not captured by standard laboratory-based evolution. This is particularly relevant when using experimental evolution as a tool to exploit trade-offs associated with resistance acquisition. Epistatic interactions are important to consider, as the phenotypic effect of a mutation may differ depending on genetic background(14). Trade-offs such as collateral sensitivity or reduced fitness may be present in one environment but absent in another. Therefore, it is important that mutational landscapes derived from adaptive laboratory evolution are comparable to those observed *in vivo*.

Our findings demonstrate that key within-patient resistance mechanisms can be reproduced within laboratory-based evolution experiments, but that resistance phenotypes and broader mutational landscapes observed *in vitro* are influenced by population bottleneck size. This suggests that stringent bottlenecks may be useful when designing adaptive laboratory evolution experiments intended to model infections where naturally occurring bottlenecks are expected to be small. This effect of experimental conditions should therefore be considered when designing adaptive laboratory evolution experiments aimed at informing clinical decision making.

## Methods

### Ethics statement

Ethical approval for linking the anonymised data with bacterial isolates was granted by the University of Liverpool as part of a separate study (REC reference 255669).

### Bacterial isolates

Bacterial isolates were identified from a database of stored clinical specimens and associated antibiogram data from Alder Hey Children’s NHS Foundation Trust, Liverpool, UK. This database was searched for the following criteria: blood culture isolates, identified as the same species, cultured from the same patient at multiple time points, and where a later isolate demonstrated increase in resistance to any antimicrobial compared to previous isolates. Using this search criteria, three clinical isolates of *Klebsiella grimontii* from a single patient were identified: VS1520, VS1543 and MB2616. Clinical data, including antimicrobial exposure between blood culture collection, was extracted and anonymised from the patient record. Isolates were grown on LB agar at 37 **°**C for 18 h followed by growth in LB broth (both Sigma, UK) at 37**°**C for 18 h at 200 rpm unless otherwise stated. Stocks were made using LB broth plus 20% glycerol (v/v, Sigma, UK) and stored at −80°C.

### Antimicrobial susceptibility testing

Antimicrobial susceptibility testing was performed for the three clinical isolates (VS1520, VS1543, and MB2616) with piperacillin sodium salt (Sigma, UK): tazobactam sodium salt (Cayman Chemical, USA) (TZP), cefuroxime sodium salt (CXM), amoxicillin trihydrate: potassium clavulanate (4:1, AMC), cefpodoxime (CPD), aztreonam (ATM), and ciprofloxacin (all Sigma, UK), using the broth microdilution method described in the EUCAST guidelines(33, 34). For TZP, piperacillin was serially diluted and tazobactam was fixed at 4 µg/mL. Antimicrobial susceptibility testing for cefoxitin (FOX) and meropenem (MEM) (both Oxoid, UK) were determined by disk diffusion method(35). TZP broth microdilution for *in vitro*-evolved isolates (*n* = 44) was performed as above. All assays were performed with three technical and three biological replicates.

Collateral susceptibility testing was performed on a subset of isolates (*n* = 8) against colistin, gentamicin, ciprofloxacin, meropenem, tigecycline, and trimethoprim-sulfamethoxazole using broth microdilution for colistin and disk diffusion for all other antibiotics.

Piperacillin sodium salt, tazobactam sodium salt, cefuroxime sodium salt, amoxicillin trihydrate: potassium clavulanate, aztreonam, and colistin were solubilised in molecular-grade water (Sigma, UK). Cefpodoxime was solubilised in molecular-grade ethanol (Sigma, UK), and ciprofloxacin in 0.1 N hydrochloric acid solution (Sigma, UK). Antibiotics were filter-sterilised using a 0.22 µM polyethersulfone (PES) membrane filter (Millipore, USA). Antimicrobial resistance phenotypes were interpreted according to EUCAST clinical breakpoints (v14.0)(34).

### *In vitro* selection for AMR

The MIC of TZP against the susceptible ancestor VS1520 was determined as 1/4 µg/mL, with TZP concentrations reported in the format piperacillin/tazobactam (µg/mL). To investigate TZP resistance evolution under different selective pressures, adaptive laboratory evolution was performed under three conditions: (i) LB with a 0.1% bottleneck (low passage volume), (ii) LB with a 5% bottleneck (higher passage volume), and (iii) LB supplemented with 5% sheep blood with a 0.1% bottleneck. The susceptible ancestor VS1520 was grown on LB agar at 37°C for 18 h, then a sweep of colonies were inoculated in LB broth (Sigma, UK) containing sub-inhibitory concentrations of TZP (0.5/0.0625 µg/mL) at 37°C for 18 h at 200 rpm, followed by either 10 µL (0.1% bottleneck) or 500 µL (5% bottleneck) culture passaged to fresh 10 mL LB broth supplemented with inhibitory concentrations of TZP (1/0.125 µg/mL). After 18 h at 37°C and 200 rpm, 50 µL of neat culture was plated onto LB agar containing 4/0.5 µg/mL and 8/1 µg/mL TZP and incubated at 37°C for 18 h. Resistant colonies were counted and three were selected from each plate and banked in LB plus 20% glycerol (v/v) at −80°C. The same experiment was performed using a 10 µL bottleneck volume (0.1%) but with the addition of defibrinated sheep blood (5% v/v) to all media (E&O Laboratories Ltd, Scotland, UK) to assess the impact of environment while controlling for bottleneck size. Sheep blood was used as a representative blood-supplemented medium to better replicate aspects of the host environment and allow comparisons between *in vitro*-evolved mutants and patient-derived bloodstream isolates. Antibiotic-free controls were run in parallel in all experimental evolution groups using the same methodology but with non-selective LB media, with one colony banked per biological replicate. The bottleneck size corresponds to the volume of bacterial culture passaged each day. Laboratory-evolved isolate IDs encode the TZP selection concentration, bottleneck volume, biological replicate, and isolate number. Isolates evolved in LB + 5% sheep blood are denoted by [B]. For example, 4 (10) 1.1 [B] indicates an isolate selected on 4/0.5 µg/mL TZP under a 0.1% bottleneck in blood-supplemented media from biological replicate 1, isolate 1. Each experiment was performed with three technical and five biological replicates, as two out of five replicates did not yield any resistant colonies. TZP ratio used was 8:1 throughout *in vitro* evolution experiments to mimic clinical dosage.

### Nitrocefin assay

A nitrocefin assay was performed to compare the β-lactamase activity of the *K. grimontii* clinical isolates (*n = 3*) and the laboratory-evolved strains (*n = 44*). Following 18 h of growth at 37°C on LB agar, single colonies were inoculated in 10 mL LB broth and incubated at 37°C for 18 h shaking at 200 rpm. The following day, cultures were diluted with fresh LB broth to an absorbance at 600 nm of 0.3 using Jenway 7205 UV/Vis spectrophotometer (Cole-Parmer Jenway, UK). Nitrocefin (Oxoid, UK) was added to each culture (10 µL) in a 96-well plate to a final concentration of 50 µM. Hydrolysis of the chromogenic β-lactam substrate nitrocefin was immediately quantified over 1 hr by measuring absorbance at 486 nm every 90 s using a Clariostar microplate reader (BMG Labtech, Germany). A blank consisting of LB broth containing 50 µM nitrocefin was included to account for background substrate hydrolysis and the ancestral strain VS1520 was included as a reference for comparison of relative β-lactamase activity. Repeats were performed in biological and technical triplicate (*n* = 3), and standard error of the mean (SEM) determined. In the clinical isolates, nitrocefin hydrolysis was calculated as mean absorbance at 486 nm over 60 min and data reported as mean ± SEM with raw replicate values provided in Supplementary Table 1. Relative nitrocefin activity of LB-evolved and LB + 5% sheep blood-evolved isolates was assessed by calculating the area under the curve (AUC) after 30 min using GraphPad Prism (v10.2.3). The AUC for each evolved isolate was then normalised by dividing AUC of the ancestor to generate a relative activity value. Results were visualised using GraphPad Prism (v10.2.3).

### Comparative fitness

The relative fitness of *in vitro*-evolved isolates was assessed in their respective environments: LB for LB (0.1% bottleneck) and LB (5% bottleneck) conditions, and LB supplemented with 5% defibrinated sheep blood for LB + blood (0.1% bottleneck) conditions. Overnight cultures were adjusted to an OD_600_ of 0.1 in LB and diluted 10^-3^ before inoculating 150 µL into flat-bottom 96-well plates. Growth was monitored using a Clariostar microplate reader at 37°C with orbital shaking at 200 rpm, measuring absorbance every 10 min for 24 h in LB, or 12 h in LB + 5% sheep blood. Absorbance in LB was measured at 600 nm, and at 615 nm for blood-supplemented media to minimise interference from haemoglobin absorption(36). All assays were performed with three technical and three biological replicates.

Relative fitness of each isolate was calculated using area under curve (AUC) values obtained in GraphPad Prism (v10.2.3), normalised to the ancestral strain VS1520 as shown below:

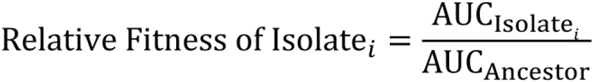

### Whole-genome sequencing (WGS)

*Short-read sequencing*: All isolates were whole-genome sequenced by MicrobesNG (http://microbesNG.com) according to their methods (v20241308). Briefly, genomic DNA libraries were prepared using the Nextera XT Library Prep Kit (Illumina, San Diego, USA), and short-read sequencing was performed using NovaSeq 6000 Illumina platform (2 x 250 bp paired-end reads). MicrobesNG also carried out adaptor trimming and quality filtering with Trimmomatic (v0.30)(37) with a sliding window quality cutoff of Q15.

*Long-read sequencing of clinical isolates VS1520, VS1543, and MB2616*: Genomic DNA was extracted using Fire Monkey High Molecular Weight DNA Extraction Kit (RevoluGen, UK) following overnight growth in LB broth at 37°C for 18 h at 200 rpm. Long-read sequencing of the three clinical isolates was performed using an R9.4.1 flow cell on a MinION sequencer (Oxford Nanopore Technologies, UK). The reads were base called using the super accuracy model and demultiplexed using Guppy (v6.4.6).

### Bioinformatics

*De novo assembly of clinical isolates VS1520, VS1543, and MB2616*: Long-read sequences were trimmed using Porechop (v0.2.4; https://github.com/rrwick/Porechop) filtered using Filtlong (v0.2.1; https://github.com/rrwick/Filtlong) with a minimum read length of 1 kb and a kept-base percentage of 90%. Long-read sequences were assembled using Flye (v2.9.2-b1786)(38), and the draft assembly was polished with long reads using Medaka (v1.5.0; https://github.com/nanoporetech/medaka). The assembly then underwent short-read polishing with Polypolish (v0.5.0)(39), followed by another round of short-read polishing using Pypolca (v0.3.1)(40). Assembled genomes were annotated using RAST annotation server(41) and visualised with SnapGene Viewer (v7.2.1). Paired short-reads of VS1543 and MB2616, and laboratory-evolved isolates (*n* = 44), were compared to a hybrid assembled genome of ancestral VS1520 using Breseq (v0.38.3)(42) to identify any accumulated SNPs and indels. Consensus mode (default) was run, which reports SNPs when at least 80% of the reads at a given position support the variant. Heatmaps were visualised with R package Pheatmap (v1.0.12; https://github.com/raivokolde/pheatmap) using R version 4.4.1 (2024-06-14 ucrt). Default methods were used for all programmes unless otherwise specified.

### Species identification and AMR genes

Species identification and multi-locus sequence type was determined using Pathogenwatch (v22.3.8; https://pathogen.watch). Average nucleotide identity (ANI) was calculated between clinical isolates using pyani (v0.2.11) with the ANIm method, which uses whole-genome alignment with MUMmer(43). Antimicrobial resistance genes were identified using ResFinder (v4.6.0)(44). Plasmids were detected with PlasmidFinder (v2.1)(45). Comparison of *K. grimontii bla*_OXY-6-4_ β-lactamase gene promoter sequences in within-patient evolved and lab-evolved strains were performed using Clustal Omega(46) and visualised with SnapGene Viewer (v7.2.1).

### Statistical analysis

A one-way analysis of variance (ANOVA) was performed to test the differences in mean nitrocefin hydrolysis between clinical strains (VS1520, VS1543, and MB2616). Tukey’s multiple comparisons test was used as a post hoc analysis to determine pairwise differences. Data were reported as mean ± standard error of the mean (SEM). To compare both nitrocefin hydrolysis (AUC), and relative fitness (AUC) between the ancestral strain (VS1520) and the evolved strains, a one-way ANOVA was performed to assess the differences among the groups. Post hoc comparisons were carried out using Dunnett’s test to specifically compare each evolved strain to the ancestral control (VS1520). Statistical significance was set at *p* < 0.05. All analyses were performed using GraphPad Prism (v10.2.3).

## Data availability

Adaptor-trimmed and quality-filtered Illumina short-read sequencing data of all isolates, together with trimmed Oxford Nanopore long-read sequencing data of the three clinical isolates, have been deposited in the NCBI Sequence Read Archive (SRA) under BioProject accession PRJNA1292152.

## Author contributions

Conceptualisation: E.A., A.P.R. and F.E.G.; formal analysis: E.A., R.N.G., A.P.R. and F.E.G.; experimental work: E.A.; resources: C.M.P., A.K., E.D.C. and R.M.; supervision: A.P.R., F.E.G., C.M.P. and N.A.F.; funding acquisition: A.P.R., E.D.C. and N.A.F.; writing—original draft: E.A.; writing—review and editing: all authors.

